# Increased actin binding is a shared molecular consequence of numerous spinocerebellar ataxia mutations in β-III-spectrin

**DOI:** 10.1101/2023.02.20.529285

**Authors:** Alexandra E. Atang, Amanda R. Keller, Sarah A. Denha, Adam W. Avery

## Abstract

Spinocerebellar ataxia type 5 (SCA5) is a neurodegenerative disease caused by mutations in the *SPTBN2* gene encoding the cytoskeletal protein β-III-spectrin. Previously, we demonstrated that a L253P missense mutation, localizing to the β-III-spectrin actin-binding domain (ABD), causes increased actin-binding affinity. Here we investigate the molecular consequences of nine additional ABD-localized, SCA5 missense mutations: V58M, K61E, T62I, K65E, F160C, D255G, T271I, Y272H, and H278R. We show that all of the mutations, similar to L253P, are positioned at or near the interface of the two calponin homology subdomains (CH1 and CH2) comprising the ABD. Using biochemical and biophysical approaches, we demonstrate that the mutant ABD proteins can attain a well-folded state. However, thermal denaturation studies show that all nine mutations are destabilizing, suggesting a structural disruption at the CH1-CH2 interface. Importantly, all nine mutations cause increased actin binding. The mutant actin-binding affinities vary greatly, and none of the nine mutations increase actin-binding affinity as much as L253P. ABD mutations causing high-affinity actin binding, with the notable exception of L253P, appear to be associated with early age of symptom onset. Altogether, the data indicate increased actin-binding affinity is a shared molecular consequence of numerous SCA5 mutations, which has important therapeutic implications.

## Introduction

Spinocerebellar ataxia type 5 (SCA5) is a neurodegenerative disease stemming from autosomal dominant mutations in the *SPTBN2* gene encoding the cytoskeletal protein β-III-spectrin. β-III-spectrin is highly expressed in cerebellar Purkinje cells^1^, the cell population targeted in SCA5 pathogenesis^2^. Within Purkinje cells, β-III-spectrin localizes to the soma and dendrites^1,3^. Mouse and cultured neuron models showed that β-III-spectrin function is required for normal dendritic arborization, spine morphology, and synaptic signaling^3-7^. Structurally, β-III-spectrin contains an N-terminal actin-binding domain (ABD), followed by seventeen spectrin-repeat domains, and a C-terminal pleckstrin homology domain. Analogous to other β-spectrins, the functional unit of β-III-spectrin is considered a heterotetramer composed of two β-III-spectrin subunits and two α-II-spectrin subunits. Super-resolution imaging of cortically localized β-III-spectrin supports a role for the β-III/α-II-spectrin heterotetramer in crosslinking actin filaments beneath the plasma membrane^7,8^. This spectrin-actin network, together with the spectrin adaptor protein, ankyrin-R, is likely important for maintaining proper subcellular localization of post-synaptic membrane proteins^3,9,10^. Many SCA5 mutations are known to localize to the ABD of β-III-spectrin^11^, supporting the importance of actin binding and the formation of a spectrin-actin network to β-III-spectrin function.

Initial mapping of three SCA5 mutations to the *SPTBN2* gene was performed in 2006^2^. These mutations, which included an ABD-localized L253P missense mutation, are associated with atrophy of the cerebellar cortex, and adult onset of symptoms that included progressive limb and gait ataxia, slurred speech and abnormal eye movements^12-14^. More recently, additional heterozygous, dominant *SPTBN2* missense mutations were reported that are associated with early onset (infantile to adolescent) of symptoms. In addition to cerebellar atrophy and ataxia, these early onset cases are characterized by delayed motor development and sometimes intellectual disability^11,15-17^. These more severe symptoms resemble those of the neurodevelopmental disorder SPARCA1^18^/SCAR14^19^, caused by homozygous recessive *SPTBN2* mutations. The SPARCA1/SCAR14 mutations are truncating mutations that are typically loss-of-function associated. Significantly, clinically normal parents heterozygous for the SPARCA1/SCAR14 loss-of-function alleles point away from haploinsufficiency as a disease mechanism for the heterozygous SCA5 mutations. For SCA5 mutations, the underlying dominant negative molecular mechanisms responsible for variability in age of onset and symptom severity are not known. Indeed, most SCA5 mutations have not been experimentally investigated at the molecular or cellular level.

We previously characterized the ABD-localized L253P missense mutation. The native L253 residue is positioned at the interface of the two calponin-homology subdomains (CH1 and CH2) comprising the ABD. Biochemically we showed that the L253P mutation causes a ∼1000-fold increase in actin-binding affinity^20^. Through biophysical approaches, including cryo-EM, we showed that high-affinity actin binding caused by the CH2-localized L253P mutation, is due to an opening of the CH1-CH2 interface^21^. This alleviates CH2 occlusion of CH1, allowing CH1 to bind actin. We further showed that L253P induced high-affinity actin binding also requires an N-terminal helix, preceding CH1, that directly binds actin^22^. In neurons, L253P reduces dendritic localization of β-III-spectrin, and induces dendritic arbor defects in cultured Purkinje cells^7^ and Drosophila sensory neurons^23^. Truncation of the N-terminal helix preceding CH1 abolishes L253P-induced high-affinity actin binding and rescues L253P-induced dendritic arbor defects and neurotoxicity in Drosophila^22^, supporting aberrant actin binding as a pathogenic mechanism. How other ABD-localized SCA5 mutations impact the structure and function of the ABD has yet to be explored.

In the present work, we characterize the molecular consequences of nine additional, heterozygous, missense mutations localizing to the β-III-spectrin ABD. Unlike L253P, many of these mutations are associated with early onset symptoms.

## Results

### Clinical summary of ABD-localized mutations

Here we report the molecular characterization of nine ataxia-associated, ABD-localized, missense mutations in β-III-spectrin. These mutations include: V58M, K61E, T62I, K65E, F160C, D255G, T271I, Y272H, and H278R. All mutations were heterozygous in patients, consistent with a dominant mechanism of action. T62I, K65E, F160C, D255G, and T271I were identified in patients with infantile onset (<12 mo) symptoms^11,15,24,25^. The compound heterozygous mutation Y272H/W2065* was identified in an infantile-onset patient^25^. Notably, parents carrying a single mutated allele (Y272H/+ or W2065*/+) were clinically normal (age of parents not reported). Thus, Y272H appears to genetically interact with W2065* to cause disease. This suggests that Y272H by itself is either insufficiently disruptive to cause symptoms, or the mutation causes mild, late onset symptoms not yet presenting in the Y272H/+ parent. For this study, only the normal clinical phenotype of the heterozygous (Y272H/+) parent will be considered. H278R was associated with adolescent symptom onset (11 years)^26^. K61E was early onset at an unconfirmed age, but evaluated at 9 years^24^. V58M was associated with adult onset (31 years)^24^. In addition to T271I and T62I characterized in this study, T271N^27^ and T62N^28^ mutations, not characterized here, are also associated with early age of onset. A detailed summary of clinical features is provided in Table 1.

**Table 1.**
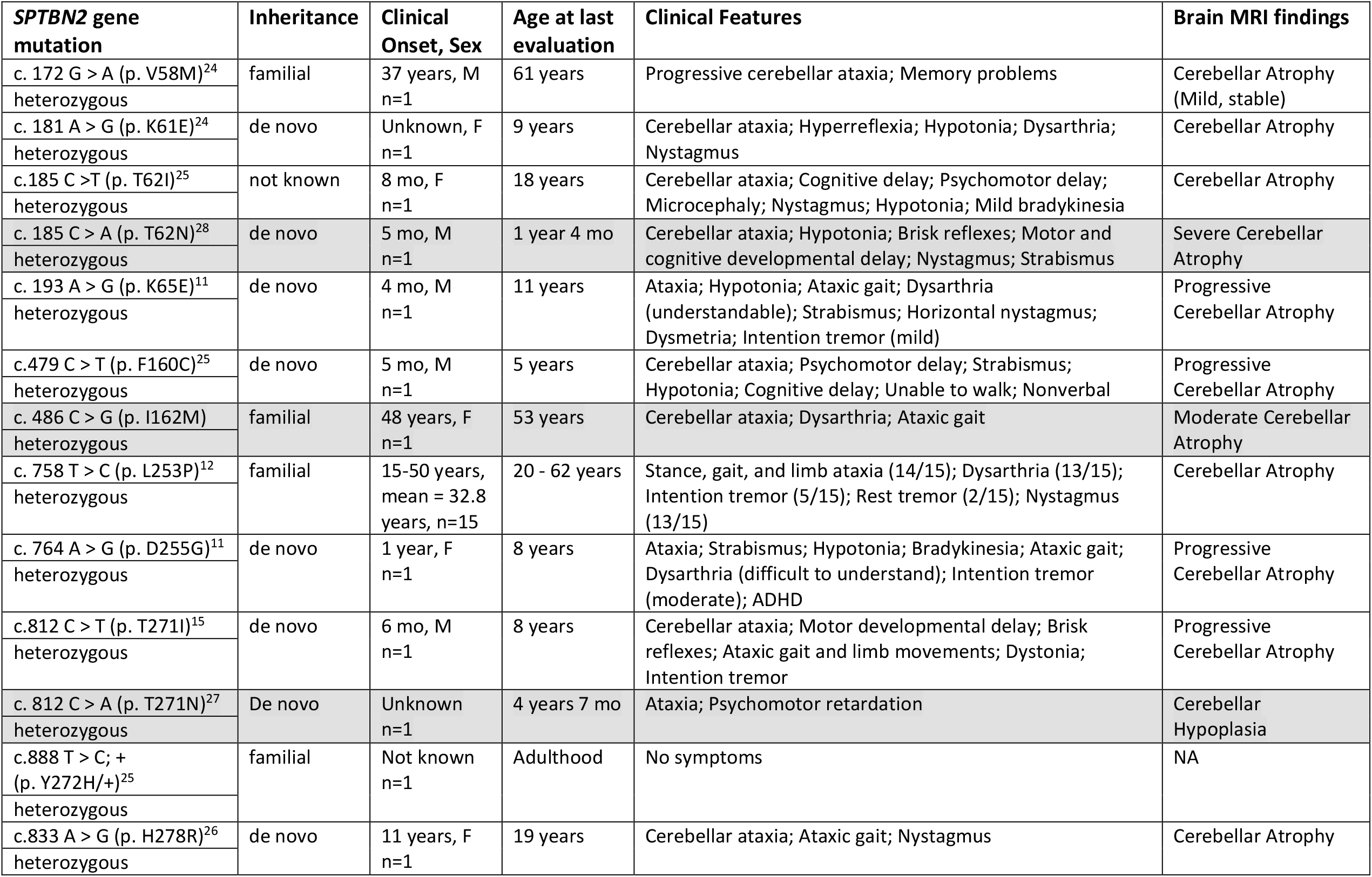
Clinical presentations of ataxia-associated mutations in the actin-binding domain of β-III-spectrin. Mutations not characterized in this study are shaded in grey.

| **SPTBN2 gene mutation** | **Inheritance** | **Clinical Onset, Sex** | **Age at last evaluation** | **Clinical Features** | **Brain MRI findings** |
| --- | --- | --- | --- | --- | --- |
| c. 172 G > A (p. V58M)<sup>24</sup><br>heterozygous | familial | 37 years, M<br>n=1 | 61 years | Progressive cerebellar ataxia; Memory problems | Cerebellar Atrophy (Mild, stable) |
| c. 181 A > G (p. K61E)<sup>24</sup><br>heterozygous | de novo | Unknown, F<br>n=1 | 9 years | Cerebellar ataxia; Hyperreflexia; Hypotonia; Dysarthria; Nystagmus | Cerebellar Atrophy |
| c.185 C > T (p. T62I)<sup>25</sup><br>heterozygous | not known | 8 mo, F<br>n=1 | 18 years | Cerebellar ataxia; Cognitive delay; Psychomotor delay; Microcephaly; Nystagmus; Hypotonia; Mild bradykinesia | Cerebellar Atrophy |
| c. 185 C > A (p. T62N)<sup>28</sup><br>heterozygous | de novo | 5 mo, M<br>n=1 | 1 year 4 mo | Cerebellar ataxia; Hypotonia; Brisk reflexes; Motor and cognitive developmental delay; Nystagmus; Strabismus | Severe Cerebellar Atrophy |
| c. 193 A > G (p. K65E)<sup>11</sup><br>heterozygous | de novo | 4 mo, M<br>n=1 | 11 years | Ataxia; Hypotonia; Ataxic gait; Dysarthria (understandable); Strabismus; Horizontal nystagmus; Dysmetria; Intention tremor (mild) | Progressive Cerebellar Atrophy |
| c.479 C > T (p. F160C)<sup>25</sup><br>heterozygous | de novo | 5 mo, M<br>n=1 | 5 years | Cerebellar ataxia; Psychomotor delay; Strabismus; Hypotonia; Cognitive delay; Unable to walk; Nonverbal | Progressive Cerebellar Atrophy |
| c. 486 C > G (p. I162M)<br>heterozygous | familial | 48 years, F<br>n=1 | 53 years | Cerebellar ataxia; Dysarthria; Ataxic gait | Moderate Cerebellar Atrophy |
| c. 758 T > C (p. L253P)<sup>12</sup><br>heterozygous | familial | 15-50 years, mean = 32.8 years, n=15 | 20 - 62 years | Stance, gait, and limb ataxia (14/15); Dysarthria (13/15); Intention tremor (5/15); Rest tremor (2/15); Nystagmus (13/15) | Cerebellar Atrophy |
| c. 764 A > G (p. D255G)<sup>11</sup><br>heterozygous | de novo | 1 year, F<br>n=1 | 8 years | Ataxia; Strabismus; Hypotonia; Bradykinesia; Ataxic gait; Dysarthria (difficult to understand); Intention tremor (moderate); ADHD | Progressive Cerebellar Atrophy |
| c.812 C > T (p. T271I)<sup>15</sup><br>heterozygous | de novo | 6 mo, M<br>n=1 | 8 years | Cerebellar ataxia; Motor developmental delay; Brisk reflexes; Ataxic gait and limb movements; Dystonia; Intention tremor | Progressive Cerebellar Atrophy |
| c. 812 C > A (p. T271N)<sup>27</sup><br>heterozygous | De novo | Unknown<br>n=1 | 4 years 7 mo | Ataxia; Psychomotor retardation | Cerebellar Hypoplasia |
| c.888 T > C; + (p. Y272H/+)<sup>25</sup><br>heterozygous | familial | Not known<br>n=1 | Adulthood | No symptoms | NA |
| c.833 A > G (p. H278R)<sup>26</sup><br>heterozygous | de novo | 11 years, F<br>n=1 | 19 years | Cerebellar ataxia; Ataxic gait; Nystagmus | Cerebellar Atrophy |

### ABD-localized mutations cluster to the CH1-CH2 interface

To explore the position of the mutations in the β-III-spectrin ABD, a structural homology model was generated using iTasser^29^, in which the plectin ABD (PDB ID: 1MB8) was the top template. The β-III-spectrin ABD is comprised of two subdomains, calponin homology domains 1 and 2 (CH1 and CH2), Figure 1A. Strikingly, all of the mutations are positioned at or near the interface of CH1 and CH2, like L253P. The native V58, K61, T62 and K65 amino acids localize to CH1 helix A, and are predicted to directly contact CH2 residues. Notably, T62 and K65 directly contact CH2 residue L253, interactions we previously highlighted in our characterization of the L253P mutation^20^. The native F160 residue localizes to CH1 helix F. The F160 sidechain is not directly oriented towards CH2. However, F160 is predicted to contact CH1 residue, W66, which in turn contacts CH2 residues L253 and T271. Mutations localizing to CH2 include D255G, T271I, Y272H, H278R and L253P. The native D255 localizes to a loop connecting CH2 helices E and F, and is predicted to form salt bridges with CH1 residues, K61 and K65. T271, Y272 and H278 localize to CH2 helix G. T271 is predicted to directly contact CH1 residues W66 and I157, in addition to contacting CH2 residue L253. Y272 forms extensive hydrophobic contacts with L253, suggesting Y272 plays an important role in coordinating the position of L253. In the homology model, H278, while oriented towards CH1, is not predicted to contact CH1 residues. To further evaluate the position of H278, we analyzed the equivalent amino acid (H275) in a crystal structure of the isolated CH2 subdomain of β-II-spectrin^30^. Alignment of the β-II-spectrin CH2 domain with the β-III-spectrin ABD homology model shows that β-II-spectrin H275 is oriented towards CH1 and predicted to contact CH1 Q161. Thus, this suggests that β-III-spectrin H278 also contacts CH1 Q161. In sum, the mutated residues cluster to the CH1-CH2 interface. Many of the mutated residues contact each other, forming an interaction network likely important for bridging CH1 and CH2. The common position of these mutations at the CH1-CH2 interface suggests that the mutations have similar structural and functional impacts on the ABD.

**Figure 1.**
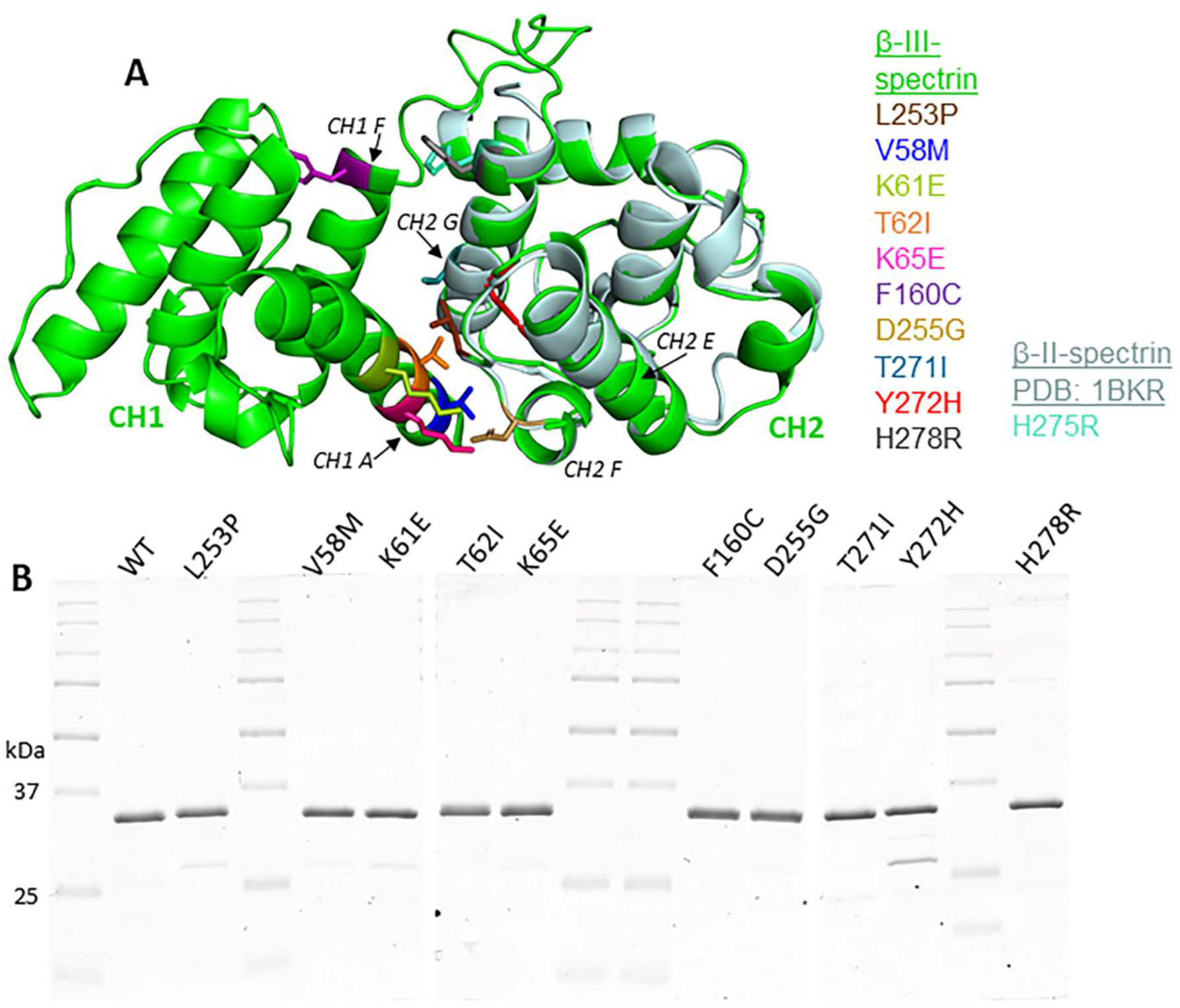
Position of SCA5 missense mutations in the actin-binding domain, and purification of mutant ABDs. **A**. Homology model of β-III-spectrin actin-binding domain (ABD; green), containing CH1 (left) and CH2 (right) subdomains. Amino acids are color-coded for sites of mutations. The crystal structure of the isolated β-II-spectrin CH2 domain (blue-silver) is aligned with β-III-spectrin CH2. All mutated residues localize to the interface of the CH1 and CH2 subdomains. **B**. Coomassie blue stained gel images showing purified wild-type (WT) and mutant ABD proteins. All ABD proteins run at the predicted size of 32 kDa.

### SCA5 mutant ABDs can attain a well-folded state, but are destabilized

To test how the SCA5 mutations impact the structure and function of the ABD, we expressed the wild-type and mutant ABD proteins in *E. coli* and performed protein purification. All ABD proteins were purified to homogeneity or contain only minor contaminating bands, Figure 1B. To determine if the SCA5 mutations impact the folded state of the ABD, circular dichroism (CD) spectroscopy was performed. Similar to our prior characterization^31^, the wild-type ABD shows an α-helical absorption profile with characteristic minima as 208 and 222 nm, Figure 2. Similarly, all ABD mutants showed pronounced α-helical profiles. From this we conclude that all the mutant ABDs can attain a well-folded state without significant disruption to secondary structure. To determine how the mutations impact the stability of the ABD, CD thermal denaturation studies were performed. The wild-type ABD unfolded in a cooperative manner, characterized by a two-state transition and a melting temperature (Tm) of 58.8°C (Figure 3), consistent with our prior report^31^. All mutant ABD proteins also displayed cooperative unfolding. However, the mutants unfolded at lower temperature than wild-type, with Tm values ranging from 51.0 to 57.9°C. An exception was L253P, which unfolded with a much lower Tm, 44.3°C, consistent with our prior characterization^31^. These denaturation studies, together with the common position of the mutations at the CH1-CH2 interface, suggest that the mutations have a shared effect to structurally destabilize CH1-CH2 interface.

**Figure 2.**
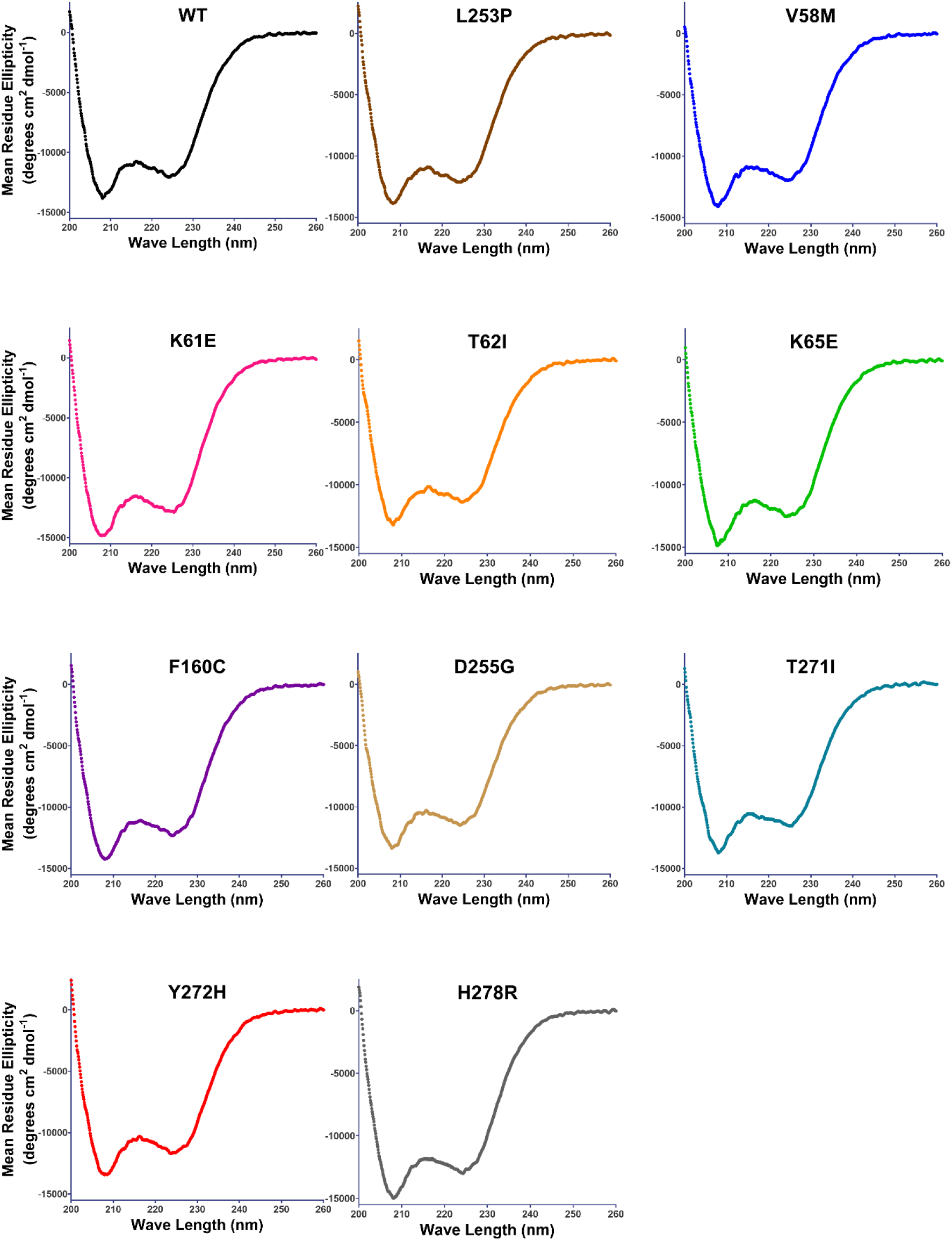
Circular dichroism absorption spectra show α-helical profiles for wild-type and mutant ABDs. CD spectra between 200 nm and 260 nm for individual wild-type, L253P, V58M, K61E, T62I, K65E, F160C, D255G, T271I, Y272H, and H278R ABD proteins. Minima at 208 and 222 nm are characteristic of α-helical fold.

**Figure 3.**
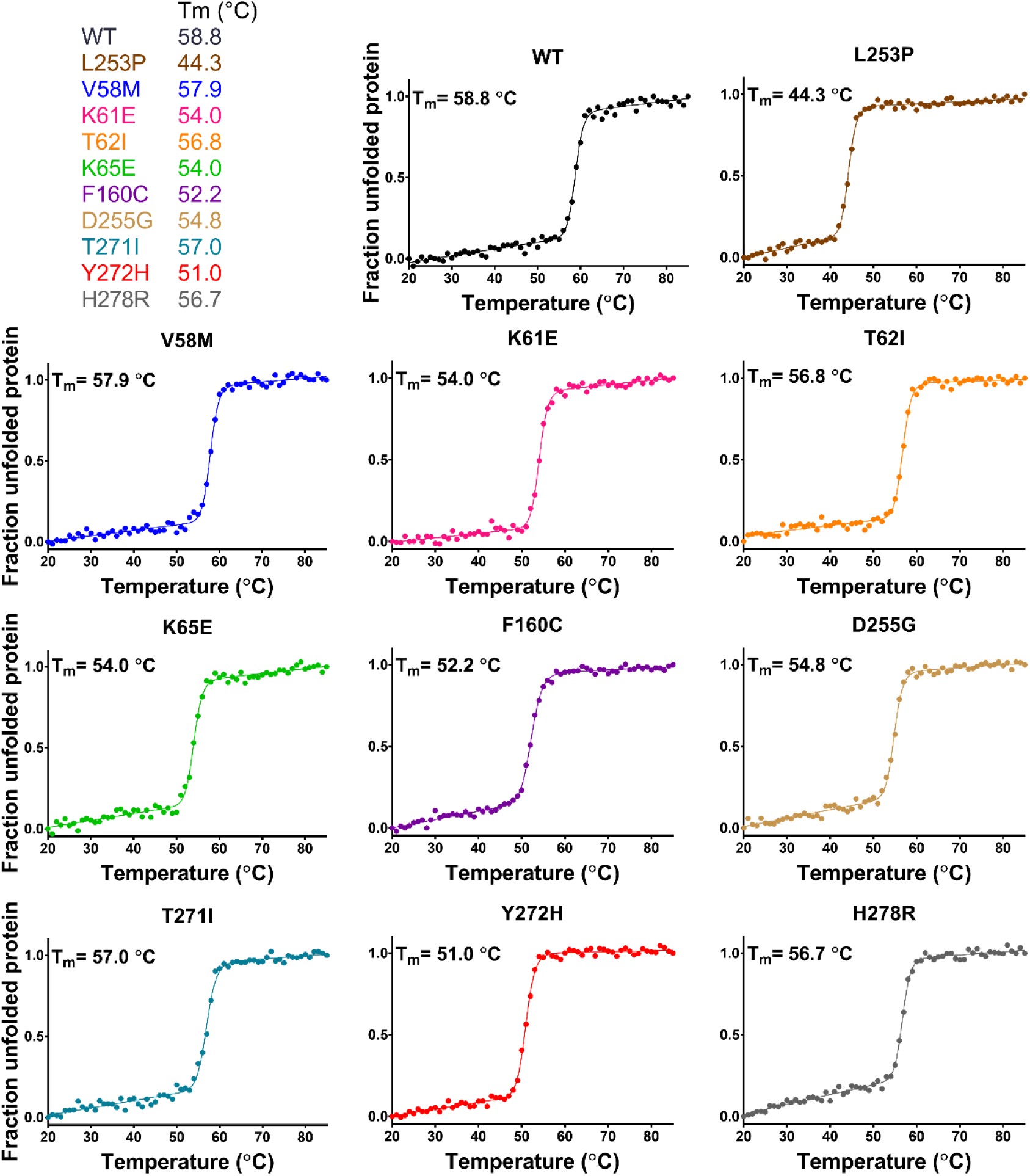
Thermal denaturation curves showing that all ABD proteins unfold cooperatively, but mutants are destabilized. Melting curves of purified ABD proteins, generated by monitoring circular dichroism absorption at 222 nm while heating sample. The cooperative, two-state, transitions indicate well-folded proteins. All mutant ABD proteins have Tm’s lower than wild-type.

### All of the SCA5 mutations increase actin-binding affinity

Previously, we showed that the L253P mutation causes a ∼1000-fold increase in actin-binding affinity, with an estimated Kd of 75 nM^31^. To determine how the additional SCA5 mutations impact actin binding, *in vitro* co-sedimentations assays were performed using the purified ABD proteins. The wild-type ABD bound F-actin with a Kd of 55.2 ± 6.7 μM, Figure 4. Significantly, all of the mutant ABDs bound actin with higher affinity than wild-type. Consistent with an estimated 75 nM Kd, the L253P ABD (2 μM) was entirely bound to F-actin at all F-actin concentrations tested (3 μM to 120 μM). Except at the lowest actin concentrations (3-10 μM), the K61E, K65E, F160C and T271I mutant ABDs were also fully bound to actin. This indicates that K61E, K65E, F160C and T271I mutations also cause high-affinity actin binding (submicromolar Kd). In contrast, V58M, T62I, D255G, Y272H and H278R mutations caused more modest increases in actin affinity. The Kd values for V58M, T62I, D255G, Y272H and H278R were 10.2 ± 1.4 μM, 2.4 ± 0.3 μM, 7.3 ± 0.5 μM, 15.3 ± 2.0 μM and 6.7 ± 1.7 μM, respectively. In sum, all of the mutations cause increased actin-binding affinity. However, the actin-binding affinities vary greatly among the different mutations, with L253P causing the highest affinity.

**Figure 4.**
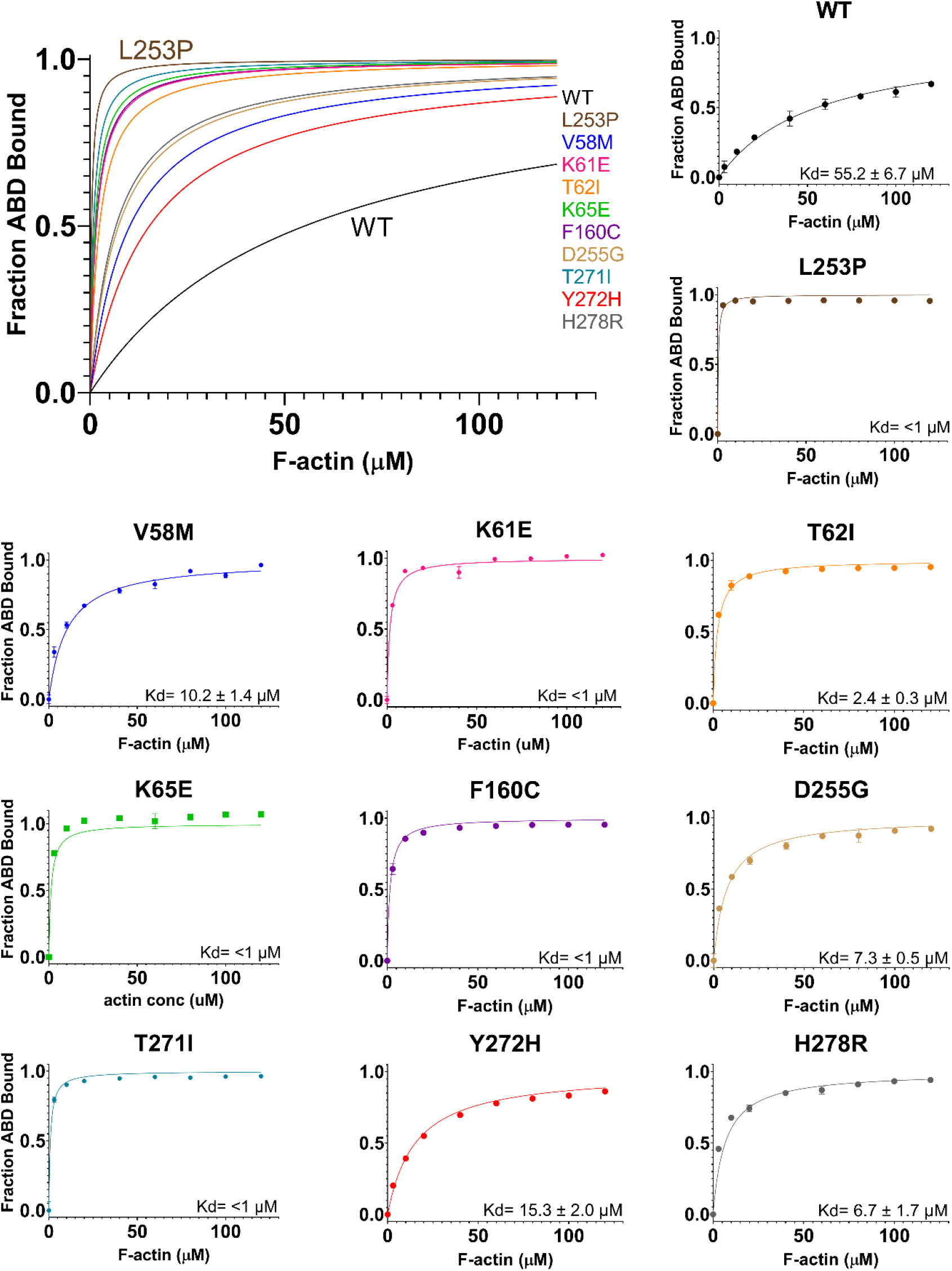
Actin co-sedimentation assays showing that all mutations increase actin-binding affinity. Co-sedimentation of ABD proteins and actin reveal greater actin affinities for all mutant ABD proteins compared to wild-type. A representative binding data and curve fit is shown for each mutant ABD protein. Kd = average +/-standard deviation, with n = 6 or more binding assays.

## Discussion

Here we build upon our prior studies of the L253P mutation by characterizing nine additional, ABD-localized missense mutations. Strikingly, our structural analyses revealed that all of the newly characterized mutations, like L253P, localize to the CH1-CH2 interface. Many of the native residues are predicted to contact each other, forming an interaction network likely important for bridging CH1 and CH2. Our circular dichroism analyses showed that the mutations are destabilizing, likely reflecting a structural uncoupling of CH1 and CH2, as we showed previously for L253P^32^. Our prior studies of the L253P mutation^32^, along with studies of spectrin-related ABD proteins^33-35^, established that opening of the CH1-CH2 interface is required for binding of CH1 to actin. Consistent with the position of the newly characterized mutations at the CH1-CH2 interface, and the destabilizing impact of these mutations, we show that all of the mutations cause increased actin binding. Thus, SCA5 mutations localizing the ABD appear to act through a common molecular mechanism.

The actin-binding affinities of the mutants vary greatly. Potentially this variability in actin-binding affinity accounts for the variability in age of onset (and symptom severity) observed for these mutations. In Table 2, our experimentally determined Kd values are listed for each mutation, along with clinical data on age of onset. With the notable exception of L253P (discussed below), mutations with submicromolar Kd values (K65E, F160C, T271I) are associated with infantile onset of symptoms. K61E also has a submicromolar Kd. However, for K61E, age of onset is not known; only age of evaluation was reported, (9y)^24^. Mutations with single digit micromolar Kd values are associated with infantile to adolescent age of onset (T62I, H278R, D255G). With a Kd value of 10 μM, V58M is associated with adult onset of symptoms. Y272H, with a Kd of 15 μM, causes the smallest increase in actin binding among the mutations. A Y272H/+ parent, of unknown age, was reported to be clinically normal^25^. Thus, the increase in actin binding caused by Y272H appears to be tolerated (no clinical phenotype) in a heterozygous state (Y272H/+). Alternatively, Y272H may cause adult onset of symptoms not yet presenting in the Y272H/+ parent. It is important to note that the nine newly characterized mutations were reported in single patients. This contrasts with the L253P mutation that was identified and evaluated in multiple (n = 15) members of a family. Age of onset for L253P ranged from 15-50y (mean = 32.8y). Thus, a correlation between actin-binding affinity and age of onset could be strengthened with higher patient number for most of the mutations.

**Table 2.** Actin-binding affinity and age of symptom onset for different ABD mutations.

| **Mutation** | **Kd (μM)** | **Age of onset** |
| --- | --- | --- |
| L253P | < 1 | 15-50 years, mean=32.8 years, n=15 |
| T271I | < 1 | 6 mo, n=1 |
| K65E | < 1 | 4 mo, n=1 |
| F160C | < 1 | 5 mo, n=1 |
| K61E | < 1 | Not reported; clinical evaluation at 9 years, n=1 |
| T62I | 2.4 ± 0.3 | 8 mo, n=1 |
| H278R | 6.7 ± 1.7 | 11 years, n=1 |
| D255G | 7.3 ± 0.5 | 12 mo, n=1 |
| V58M | 10.2 ± 1.4 | 37 years, n=1 |
| Y272H/+ | 15.3 ± 2.0 | Y272H/+ parent is clinically normal, n=1 |
| Wild-type | 55.2 ± 6.7 | NA |

Our data show that L253P is unique among the mutations in that it causes 1) the highest actin-binding affinity, and 2) the greatest protein destabilization. It thus seems likely that the decreased severity of L253P is attributed to one or both of these features. Our current and prior^31^ thermal denaturation studies show in vitro that L253P ABD begins to unfold near physiological temperature. It is thus possible that L253P β-III-spectrin is prone to denaturation and enhanced protein turnover in cells at physiological temperature. Increased protein turnover of the L253P mutant would reduce its abundance in neurons, corresponding to a reduction in neurotoxicity and a later age of symptom onset. However, in transiently transfected HEK293T cells, wild-type and L253P β-III-spectrin ABD or full-length proteins showed similar protein levels by western blot analyses^23^. Clarkson et al additionally showed that L253P β-III-spectrin did not induce pathways associated with unfolded protein response^36^.

Moreover, we showed that L253P, and a L253A mutation, which causes high-affinity actin binding, but is less destabilizing than L253P, cause similar mislocalization (absence from plasma membrane extensions and internal accumulation on actin-rich vesicles) of β-III-spectrin in HEK293T cells^23^. This supports that the behavior of the L253P mutant β-III-spectrin in cells is driven by elevated actin binding, not protein denaturation/unfolding. Alternatively, it is possible that high-affinity actin binding localizes the L253P mutant to a distinct population of actin filaments not bound by the ABD mutants with relatively lower actin-binding affinity. Future studies assessing the impact of L253P versus the other ABD mutations on mutant β-III-spectrin steady-state protein levels and subcellular localization in cultured Purkinje neurons or mouse models may clarify why L253P has reduced toxicity/late onset symptoms.

Our finding here that increased actin-binding affinity is a shared molecular consequence of numerous SCA5 mutations has important therapeutic implications. Specifically, it suggests that a small molecule drug that can alleviate high-affinity actin binding may be broadly useful as a SCA5 therapeutic. Recently we reported the identification of several small molecules capable of reducing binding of the L253P mutant to actin in vitro^37^. This established that the spectrin-actin interaction can be targeted by small molecules. Potentially these compounds will be effective in reducing actin binding caused by the multiple different SCA5 mutations characterized here. Our analysis of Kd versus age of symptom onset suggests that reducing actin-binding affinity to a Kd value of ∼15 μM (Y272H) or higher may be needed to fully eliminate toxicity caused by the ABD mutations.

## Methods

### Protein expression and purification

The following primers were used to introduce SCA5 mutations into the β-III-spectrin ABD coding sequence:

V58M forward: CTGGCAGATGAACGAGAAGCTATGCAGAAGAAAACCTTCACCAAG

V58M reverse: CTTGGTGAAGGTTTTCTTCTGCATAGCTTCTCGTTCATCTGCCAG

K61E forward: GAACGAGAAGCTGTGCAGAAGGAAACCTTCACCAAGTGGGTAAAC

K61E reverse: GTTTACCCACTTGGTGAAGGTTTCCTTCTGCACAGCTTCTCGTTC

T62I forward: GAGAAGCTGTGCAGAAGAAAATCTTCACCAAGTGGGTAAACTC

T62I reverse: GAGTTTACCCACTTGGTGAAGATTTTCTTCTGCACAGCTTCTC

K65E forward: GTGCAGAAGAAAACCTTCACCGAGTGGGTAAACTCGCACCTGGCC

K65E reverse: GGCCAGGTGCGAGTTTACCCACTCGGTGAAGGTTTTCTTCTGCAC

F160C forward: GTCTGGACCATCATCCTTCGATGCCAGATCCAAGACATCAGTGTG

F160C reverse: CACACTGATGTCTTGGATCTGGCATCGAAGGATGATGGTCCAGAC

D255G forward: GGACTTACCAAGCTGCTGGGTCCCGAAGACGTGAATGTGGACC

D255G reverse: GGTCCACATTCACGTCTTCGGGACCCAGCAGCTTGGTAAGTCC

T271I forward: GCCAGATGAGAAGTCAATCATTATCTATGTGGCTACTTACTACC

T271I reverse: GGTAGTAAGTAGCCACATAGATAATGATTGACTTCTCATCTGGC

Y272H forward: GATGAGAAGTCAATCATTACCCATGTGGCTACTTACTACCATTAC

Y272H reverse: GTAATGGTAGTAAGTAGCCACATGGGTAATGATTGACTTCTCATC

H278R forward: CTATGTGGCTACTTACTACCGTTACTTCTCCAAGATGAAG

H278R reverse: CTTCATCTTGGAGAAGTAACGGTAGTAAGTAGCCACATAG

Mutations were introduced into the β-III-spectrin ABD using the previously generated pE-SUMO-ABD WT construct^32^, through site-directed PCR mutagenesis (PfuUltra High-Fidelity DNA Polymerase, Agilent). The resulting mutant DNAs were sequence verified. pE-SUMO-ABD constructs were transformed into Rosetta 2 (DE3) *E. coli* (Novagen). 250 mL to 1 L bacteria cultures were grown in LB broth with ampicillin (100 μg/mL) and chloramphenicol (34 μg/mL). ABD protein expression was induced with 0.5 mM IPTG, for 6 h, at 300 RPM and room temperature. Bacteria cultures were pelleted at 2987 RCF for 30 min at 4 °C, and pellets stored at -20°C until further use. Bacterial pellets were resuspended in lysis buffer (50 mM Tris pH 7.5, 300 mM NaCl, 25% sucrose, and protease inhibitors (Complete Protease Inhibitor tablet, EDTA-free, Roche)), and lysed by incubation with lysozyme (Sigma) for 1 h at 4 °C, followed by a freeze-thaw cycle in an isopropanol-dry ice bath. To the lysate, MgCl_2_ (10 mM final concentration) and DNase1 (Roche) (8 U/mL final concentration) were added and incubated with slow stirring for 1 h at 4 °C. Lysates were clarified at 18,000 RPM at 4 °C for 30 min, in a Sorvall SS-34 rotor. Supernatants were syringe filtered through 0.45 μm disk filters and loaded into Poly-Prep (Biorad) chromatography columns containing Ni-NTA agarose (Qiagen) equilibrated in binding buffer containing 50 mM Tris pH 7.5, 300 mM NaCl and 20 mM imidazole. The columns were washed with binding buffer and then ABD proteins eluted with 50 mM Tris pH 7.5, 300 mM NaCl and 150 mM imidazole. For each mutant, elution fractions containing ABD protein was pooled and loaded into a Slide-a-Lyzer, 10 K MWCO, dialysis cassette (ThermoScientific) and dialysis performed overnight in 25 mM Tris pH 7.5, 150 mM NaCl, 5 mM β-mercaptoethanol, at 4 °C. 6X-His-SUMO tag was cleaved from ABD proteins using 1:10 mass ratio of Ulp1 SUMO protease:ABD in a 2 h incubation at 4 °C. Cleaved 6X-His SUMO tags and His-tagged SUMO protease were removed from the ABD proteins using 0.5 – 1 mL Ni-NTA agarose in a Poly-Prep chromatography column. Eluted fractions containing ABD proteins were pooled for each mutant and loaded into a Slide-a-Lyzer, 10 K MWCO, dialysis cassette for dialysis in 10 mM Tris pH 7.5, 150 mM NaCl, 2 mM MgCl_2_, 1 mM DTT, at 4 °C for 15 h. Recovered ABD proteins were measured for concentration using Bradford assay and snap frozen in liquid nitrogen and stored at -80 °C for future use.

### Circular dichroism measurements

Purified ABD proteins were thawed and clarified at 43,000 RPM at 4 °C for 30 min, in a Beckman TLA 100.3 rotor. Bradford assay (Biorad) was used to determine the ABD protein concentrations. ABD proteins were diluted to 150 – 200 ng/μL in buffer containing 10 mM Tris pH 7.5, 150 mM NaCl, 2 mM MgCl_2_, 1 mM DTT. CD spectra were obtained in a Jasco J-815 Spectropolarimeter with a Peltier temperature controller. A baseline correction using diluent buffer was acquired immediately before analysis. Analyses were carried out between 200 and 260 nm at 25 °C. Thermal denaturation studies for each ABD protein were obtained in a Jasco J-810 Spectropolarimeter. Unfolding was measured at 222 nm with increasing temperature from 20 – 85 °C.

### F-actin co-sedimentation assays

F-actin was prepared and purified from rabbit skeletal muscle (Pel-Freez Biologics) as previously described^22^. Purified ABD proteins were thawed and clarified at 43,000 RPM at 4 °C for 30 min, in a Beckman TLA 100.3 rotor. Bradford assay (Biorad) was used to determine the F-actin and ABD protein concentrations. As performed previously^22^, binding reactions were prepared with 2 μM ABD protein and increasing concentrations of F-actin (3 – 120 μM) in F-buffer (10 mM Tris pH 7.5, 150 mM NaCl, 0.5 mM ATP, 2 mM MgCl_2_, and 1 mM DTT). After 30 min incubation at room temperature, binding reactions were centrifuged at 50,000 RPM for 30 min at 25 °C in a TLA-100 rotor (Beckman). Promptly after centrifugation, supernatants were sampled and mixed with Laemmle sample buffer (Biorad). Collected supernatant samples were separated by SDS-PAGE and bands were visualized using Coomassie Brilliant Blue R-250 (Biorad) solution. After sufficient destaining to remove background, gels were imaged using the 680 nm channel of an Azure Sapphire imager. ABD protein band fluorescence intensities were quantified from image files using Image Studio Lite version 5.2 software. A standard curve was generated using various amounts of ABD proteins on a Coomassie stained gel to relate ABD fluorescence intensity to known ABD concentrations. The standard curves were used to convert raw fluorescence ABD signals to concentration for the binding assays. Using Prism 8 (Graphpad), dissociation constants (Kd) were determined through a nonlinear regression fit for a one-site specific binding equation with Bmax constrained to 1, as described previously^31^.

### Structural modeling

The β-III-spectrin ABD structural homology model was generated through the i-Tasser server^29^. The top template structure was the plectin ABD (PDB ID: 1MB8). Analyses of the structure, including identification of predicted contacts for different mutated residues, was performed with PyMOL v2.5.4.

## Acknowledgments

We thank Emily Oumelaz, Sophie Turcotte, Andrew Januszek, and Hannah Merrow, students in Dr. Avery’s Biochemistry laboratory course, for performing pilot studies in the class on the V58M, K61E, D255G and K65E mutant ABD proteins.

## Author contributions

AWA conceived and designed experiments. AEA, ARK and SAD acquired data and interpreted results. AEA and AWA prepared the manuscript draft. All authors have read and agreed to the published version of the manuscript.

## Funding and additional information

This work was supported by NIH grant R15NS116511 to AWA.

## Data availability

All data are contained within the manuscript.

